# Caris-ComBat-seq: Directionally Harmonizing Large-Scale RNA-seq Datasets

**DOI:** 10.1101/2025.11.26.690779

**Authors:** Laura M. Richards, Mukund Varma, Noah Spies, Nicolas Stransky, Fred P. Davis

**Author notes:** Corresponding authors Correspondence to Laura M. Richards and Fred P. Davis.

## Abstract

**Background:** Bulk RNA sequencing (RNA-seq) is an essential research and clinical diagnostics tool capable of uncovering biological insights across experimental conditions, sample types and diseases at scale. However, RNA-seq data is sensitive to batch effects introduced by technical variation. Harmonizing expression (or batch-correcting) is critical when analyzing measurements from different RNA-seq platforms. In the context of oncology, efficiently and accurately harmonizing expression at scale is important for harnessing massively large datasets (hundreds of thousands of samples) of tumor molecular profiles from different assay platforms. We aimed to develop a method for harmonizing expression in this challenging context.

**Results:** Here, we extend the widely used ComBat-seq method, as implemented in the pyComBat tool, to enable three key advances: (i) directionally adjusting counts from one batch towards a reference batch, rather than an average expression profile, (ii) separating the training and adjusting steps so that newly profiled samples not available at the time of initial model training can be harmonized, and (iii) flexibly handling outliers to improve the quality of harmonized counts. The resulting model correctly learns gene-specific differences between assay platforms and can near-instantaneously harmonize individual samples. We validated the use of the Caris-ComBat-seq tool to harmonize RNA-seq measurements on a benchmarking dataset of ∼10,000 TCGA tumor samples. Finally, we demonstrated its strength for very large datasets, by harmonizing RNA-seq data from nearly half a million tumor samples profiled by two different next-generation sequencing assays in Caris Life Science’s clinical laboratory. Source code, tutorials and manuscript data are available at: https://github.com/Caris-Life-Sciences/Caris-ComBat-seq and https://doi.org/10.5281/zenodo.17154014.

**Conclusions:** We present Caris-ComBat-seq, a new variant of the ComBat-seq algorithm designed to harmonize count-based expression data in the context of high-throughput profiling laboratories. It offers the same ability to ameliorate batch effects and retain biological signal as the original ComBat-seq, with additional operational flexibility and scaling that benefits its high-throughput application.

## BACKGROUND

Batch effects are common in RNA-sequencing (RNA-seq) data, often created by technical variation between experiments, such as differences in experimental design, sequencing platforms, reagent kits, analysis pipelines, or environmental conditions. These batch effects are critical to consider when combining RNA-seq datasets, because technically-driven batch effects can mask true biological signal and confound analysis results (1). In the context of oncology, accounting for RNA-seq batch effects is important in a clinical diagnostic setting, where it impacts analyses such as developing and applying genomic signatures and classifying tumor subtypes (2, 3). It is also essential for the broader research community to facilitate consortia studies which can involve multiple laboratories and assay platforms, as well as for integrating published consortia datasets with individual user-generated datasets (4).

Reprocessing batches of RNA-seq data with consistent pipelines, including alignment and quantification steps, is not sufficient to overcome batch effects when combining data from multiple sources (1). Although methods pioneered by the single cell transcriptomics community can integrate millions of cells across batches into the same reduced dimensionality space, they often do not correct raw count values, and instead operate on normalized expression levels (5, 6). The bioinformatics community has developed many effective statistical frameworks to address technical batch effects in bulk RNA-seq data – including both unknown sources of variation (e.g. SVAseq and RUV-seq (7, 8)) and known batch labels (e.g., specified as covariates during differential expression analysis with limma-voom, DESeq2, and EdgeR (9–12)). ComBat-seq is one of the most highly cited and widely adopted frameworks for removing known batch effects from RNA-seq data (13).

ComBat-seq uses negative binomial regression to model the read count distribution within each batch for every gene, and then subsequently maps each count in the original gene-by-sample count matrix to a new, batch-free count distribution that represents the average of each batch’s distribution. The negative binomial modeling enables ComBat-seq to maintain the integer nature of the count data after harmonizing, a necessary feature for many downstream analytical tasks (13).

To meet the demands of a high-throughput molecular laboratory continuously profiling new samples, we developed Caris-ComBat-seq, a novel adaptation of pyComBat-seq with significant performance increases. Caris-ComBat-seq uses the same statistical framework as the original algorithm, but with three major advancements. First, Caris-ComBat-seq can directionally adjust count data from one batch towards another, as opposed to an averaged expression profile. This feature enables selection of a high-quality batch for use as a “reference”. Second, by splitting the training and adjustment steps, Caris-ComBat-seq can adjust newly profiled samples without requiring these samples to be included in a newly re-trained model. In contrast, the original algorithm requires joint training and adjusting of the new samples along with previous samples, which is impractical in a clinical laboratory continually profiling new samples. Finally, Caris-ComBat-seq offers a flexible and user-defined approach for handling outliers, which can reduce their impact on harmonized expression quality. Our modifications also reduce run-time and improve scalability by allowing parallelization of jobs across multiple cores. We demonstrate these improvements on a dataset of nearly half a million samples.

## IMPLEMENTATION

Caris-ComBat-seq builds on pyComBat-seq, the Python 3 implementation of ComBat-seq as distributed by the InMoose package (https://github.com/epigenelabs/inmoose/) (14). Caris-ComBat-seq modifies pyComBat-seq to (i) directionally harmonize, (ii) separate the training and adjusting steps, and (iii) more flexibly handle outlier detection. We will first review the original ComBat-seq framework and then describe our modifications in turn.

### Original ComBat-seq model

ComBat-seq (13) models observed read counts 𝑦_g𝑖j_ as drawing from a negative binomial (NB) distribution with a gene- (𝑔), batch- (𝑖), and sample- (𝑗) specific mean and gene/batch-specific dispersion, y_gij_ ∼ NB(µ_gij_, 𝛟_gi_). The scaled count is decomposed into three components: gene-specific mean, α_g_, the impact of the biological condition of the sample on a gene-specific regression coefficient, X_j_β_g_, and the gene-specific batch effect, 𝛾_g𝑖_; this decomposition is further adjusted by the sample-specific library size, log N_j_. The variance of this distribution is a combination of the mean scaled count and a gene- and batch-specific dispersion coefficient, 𝛟_gi_.

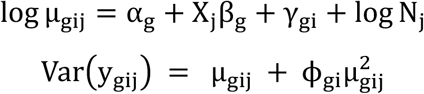

Model training infers the decomposition to build a batch-free distribution, ygj^∗^ ∼ NB(µgj^∗^, 𝛟g^∗^).

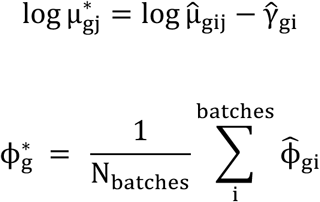

The count adjustment step then performs quantile mapping to convert values from the original distribution, 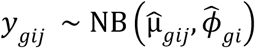, to the batch-free distribution, 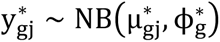.

### Directionally harmonizing

To enable directionally harmonizing towards a reference batch, we set the average expression for each gene to the value in the reference-batch, α_g_ = α_rg_, set the batch effect for the reference batch to 0, γ_gr_ = 0, and set the batch-free target dispersion to the value observed in the reference batch 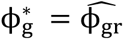. These changes make the batch-free target distribution equivalent to the reference batch’s original distribution. If a reference batch is not specified, Caris-ComBat-seq defaults to the average of batches batch-free distribution, as in the original ComBat-seq.

### Separating the training and adjusting steps

Caris-ComBat-seq separates the harmonizing process into distinct training and adjusting steps **(Figure 1A).** In the training step, we estimate for each gene the batch-specific dispersion 𝛟_gi_, batch effects γ_gi_, batch-free dispersion 𝛟_g_^∗^, and average expression α_g_. During the adjusting step, we take as input a target sample 𝑡’s observed read counts and batch label b, and then estimate for each gene the distribution mean, µ_gt_*, and its corresponding batch-free gene-specific mean, µ_gt_^∗^, given the model’s α_g_, gene-specific batch effects γ_gb_, and library size 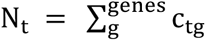. We then use these parameters to perform quantile mapping from the original batch’s 𝑁𝐵 distribution with mean µ_gt_ and dispersion 𝛟_gb_ to a batch-free distribution with mean µ_gt_^∗^ and dispersion 𝛟_g_^∗^ .

**Figure 1.**
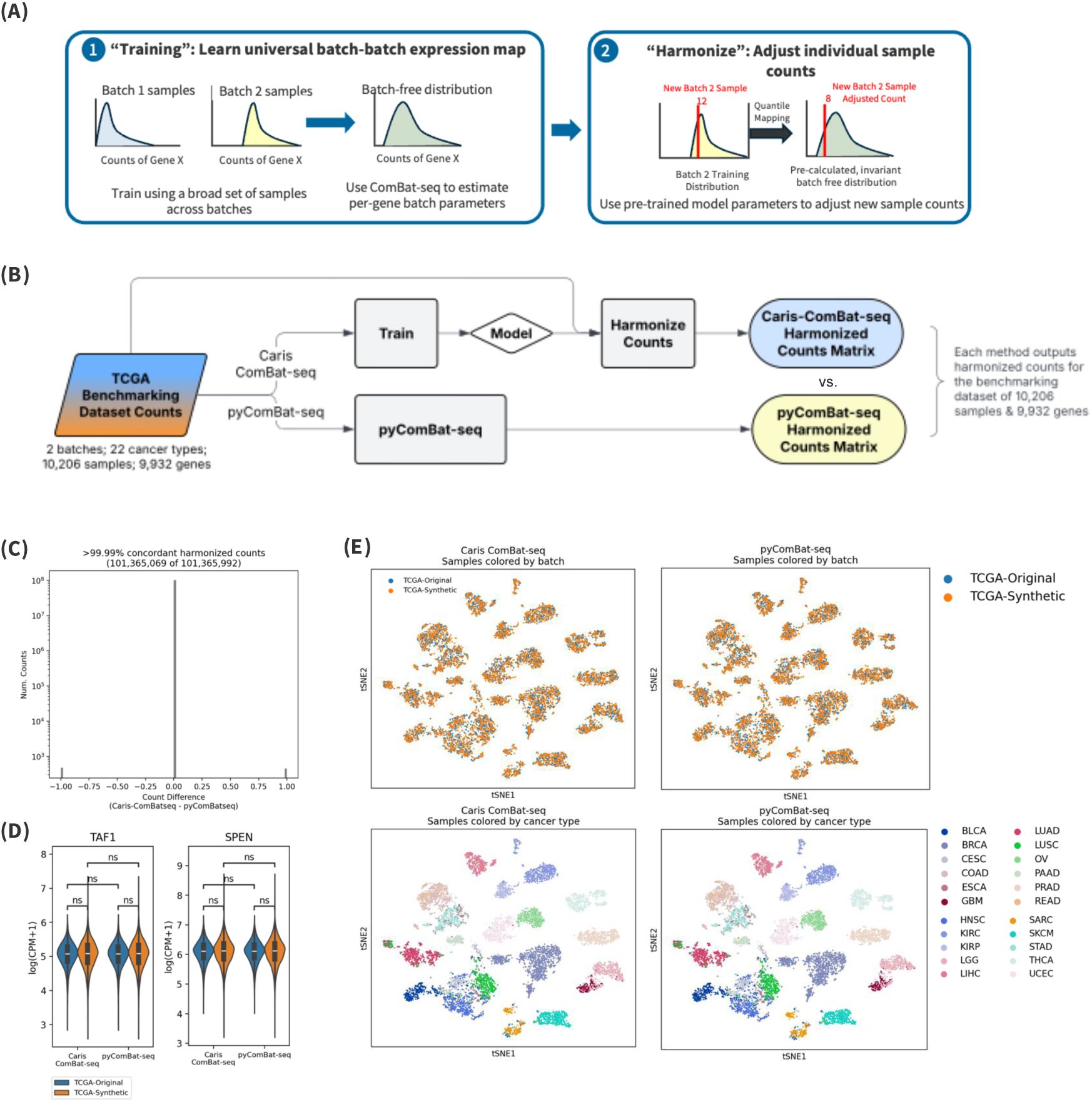
Caris-ComBat-seq recapitulates adjusted counts produced by original pyComBat-seq algorithm. **(A)** Schematic of Caris-ComBat-seq method, outlining the split of the original ComBat-seq statistical framework into two separate steps (1) training and (2) adjusting. **(B)** Flowchart detailing benchmarking experiment to compare accuracy of adjusted counts produced by default Caris-ComBat-seq to adjusted counts produced by pyComBat-seq. **(C)** Histogram depicting count difference between Caris-ComBat-seq and pyComBat-seq adjusted counts across 10,206 samples and 9,932 genes. Each point represents a single sample’s adjusted count value for a single gene. **(D)** Comparing harmonized expression levels between methods for two genes (*TAF1, SPEN*) that show significant batch effects in the raw counts data. Statistical significance of mean expression differences between batches assessed with Mann-Whitney-Wilcoxon test two-sided with Benjamini-Hochberg (BH) correction. No significance (ns): 0.05 < *P* ≤ 1.0. **(E)** *t*-SNE visualization of sample similarity using expression harmonized by Caris-ComBat-seq (left panels) and pyComBat-seq (right panels). *t*-SNEs are colored by batch (upper panels) and cancer type (lower panels). Each point represents a sample (N = 10,206).

### Outlier detection

During quantile mapping, the original ComBat-seq algorithm evaluates the likelihood of each original scaled count under the batch-specific distribution, and considers it an outlier if it is 𝑝 < 10^−4^ . If a count is an outlier, ComBat-seq does not adjust it and instead retains the original count value in the final harmonized result. We remove this hard-wired threshold and instead make it an optional parameter. If a threshold is not provided, we attempt to adjust all original counts, regardless of p-value, and only resort to retaining the original count value if the p-value is so low that quantile mapping is not technically possible (i.e. the relevant statistical routines return an Infinity value). This change reduces the number of outlier values, increasing the number of expression values that are adjusted, and improving the continuity of the adjusted expression distributions.

## RESULTS

### Benchmarking dataset and data pre-processing

Traditionally, bulk RNA-seq batch correcting algorithms have been validated on small test datasets, generally fewer than 50 samples of a single biological sample type. To validate Caris-ComBat-seq on a real-world dataset with thousands of samples and broad representation of multiple biological sample types (e.g. cancer types), we used RNA-seq data from The Cancer Genome Atlas (TCGA) (15) to create our benchmarking dataset **(Supplemental Fig. S1A)**.

To evaluate harmonized expression quality, we created two batches within the TCGA dataset, mirroring transcriptional differences one might observe between two collections of RNA-seq data generated with different assays or sequencing platforms. To accomplish this, we obtained raw gene-level counts for TCGA from the recount2 portal (16), totaling 11,837 genes in 11,284 samples across 35 cancer types. Next, we filtered this dataset to include only cancer types and protein-coding genes that are well represented in both TCGA and Caris Life Science’s internal RNA-seq database, for a final dataset size of 9,932 genes in 10,206 samples across 22 cancer types. We then split this dataset in half equally by cancer type. The first batch – termed “TCGA-Original” – was comprised of original, unmodified recount2 gene counts for 5,101 samples. To generate a second, transcriptionally distinct batch, we used Caris-ComBat-seq to directionally adjust the second half of TCGA transcriptomes (N = 5,105) to resemble samples sequenced on Caris’ clinical platform. This second batch is termed “TCGA-Synthetic” **(Supplemental Fig. S1A-B).**

As desired, we observed significantly different per-gene expression distributions between TCGA-Original and TCGA-Synthetic, confirming that the generated datasets had a batch effect suitable for benchmarking **(Supplemental Fig. S1C).** We next used *t*-SNE embedding to visualize sample mixing between batches at a transcriptome-wide level. RNA-seq counts per sample were library-size normalized to counts per million (CPM) and log transformed. Using Scanpy (17) (v1.10.3), the top 2,500 highly variable genes were identified, followed by principal component analysis (PCA) for dimension reduction and *t*-SNE using the top 20 principal components (PCs). All plotting and statistical analyses were performed with the following Python packages: Matplotlib (18) (v3.9.2), Seaborn (19) (v0.13.2), statannotations (v0.7.1; https://github.com/trevismd/statannotations) and SciPy (20) (v1.14.1). This exercise demonstrated that samples grouped strongly by batch instead of biological signal (such as cancer type), confirming the successful generation of technical artifact in the data **(Supplemental Fig. S1D-E)**.

### Caris-ComBat-seq recapitulates original pyComBat-seq results

To demonstrate that the underlying statistical framework of Caris-ComBat-seq is identical to that of pyComBat-seq, we harmonized our TCGA benchmarking dataset using both methods **(Fig. 1B).** To more accurately recapitulate the results of pyComBat-seq, we used Caris-ComBat-seq with all default parameters: (i) we first trained a model on the benchmarking dataset without a reference batch specified, and then (ii) harmonized the count values on the same input matrix used for training with pyComBat-seq’s outlier threshold of 𝑝 < 10^−4^.

Harmonized count matrices produced by pyComBat-seq and Caris-ComBat-seq were >99.99% concordant **(Fig. 1C)**. 101,365,069 of 101,365,992 individual gene-by-sample datapoints were identical between the two methods. The 923 discordant values differed by a magnitude of only one count and are likely related to float precision or rounding of the adjusted counts during quantile mapping. To further illustrate their concordant performance, we find that both tools successfully harmonized the expression levels (**Fig. 1D**) of two genes with very significant batch effects (*TAF1* and *SPEN*; **Fig. S1C**). Unlike their original counts (**Fig. S1C**), their harmonized counts had no significant difference between batches (*P*>0.05; Mann-Whitney-Wilcoxon test two-sided with BH correction; **Fig. 1D**).

Next, we assessed the integrity of the biological signal in the harmonized dataset using *t*-SNE embedding. In stark contrast to raw counts **(Fig. S1D-E),** harmonized counts produced by both pyComBat-seq and Caris-ComBat-seq did not display batch effects **(Fig. 1E).** Samples no longer grouped by technical batch (i.e. TCGA-Original vs. TCGA-Synthetic), but instead grouped by cancer type. These results suggest that biologically relevant signal dominated over technical aspects of the dataset, making it suitable for downstream analyses such as differential gene expression.

Together, these results demonstrate that Caris-ComBat-seq’s new features do not hinder its ability to recapitulate results from the original, gold-standard ComBat-seq algorithm.

### Directionally harmonizing towards a reference batch profile

Inspired by ComBat-ref, a recently developed modification of ComBat-seq (21), Caris-ComBat-seq can directionally adjust towards a single reference batch, rather than the average of all batches. To do so, a reference batch must be specified during the training step of Caris-ComBat-seq. With this approach, the counts for samples belonging to the reference batch remain unmodified and only counts from non-reference batch samples are adjusted. Using our benchmarking dataset, we demonstrate successfully harmonizing TCGA-Original counts towards the TCGA-Synthetic batch as the fixed reference, and vice versa **(Fig. 2A)**. We used *TAF1* gene expression as an example to showcase the effect of directionally harmonizing. When TCGA-Original is the reference batch, harmonized *TAF1* expression in TCGA-Original samples remain identical to their raw counts, while counts from TCGA-Synthetic samples are adjusted to resemble samples in TCGA-Original (*P*>0.05; Mann-Whitney-Wilcoxon test two-sided with BH correction). The reverse pattern is true when using TCGA-Synthetic as the reference batch.

**Figure 2.**
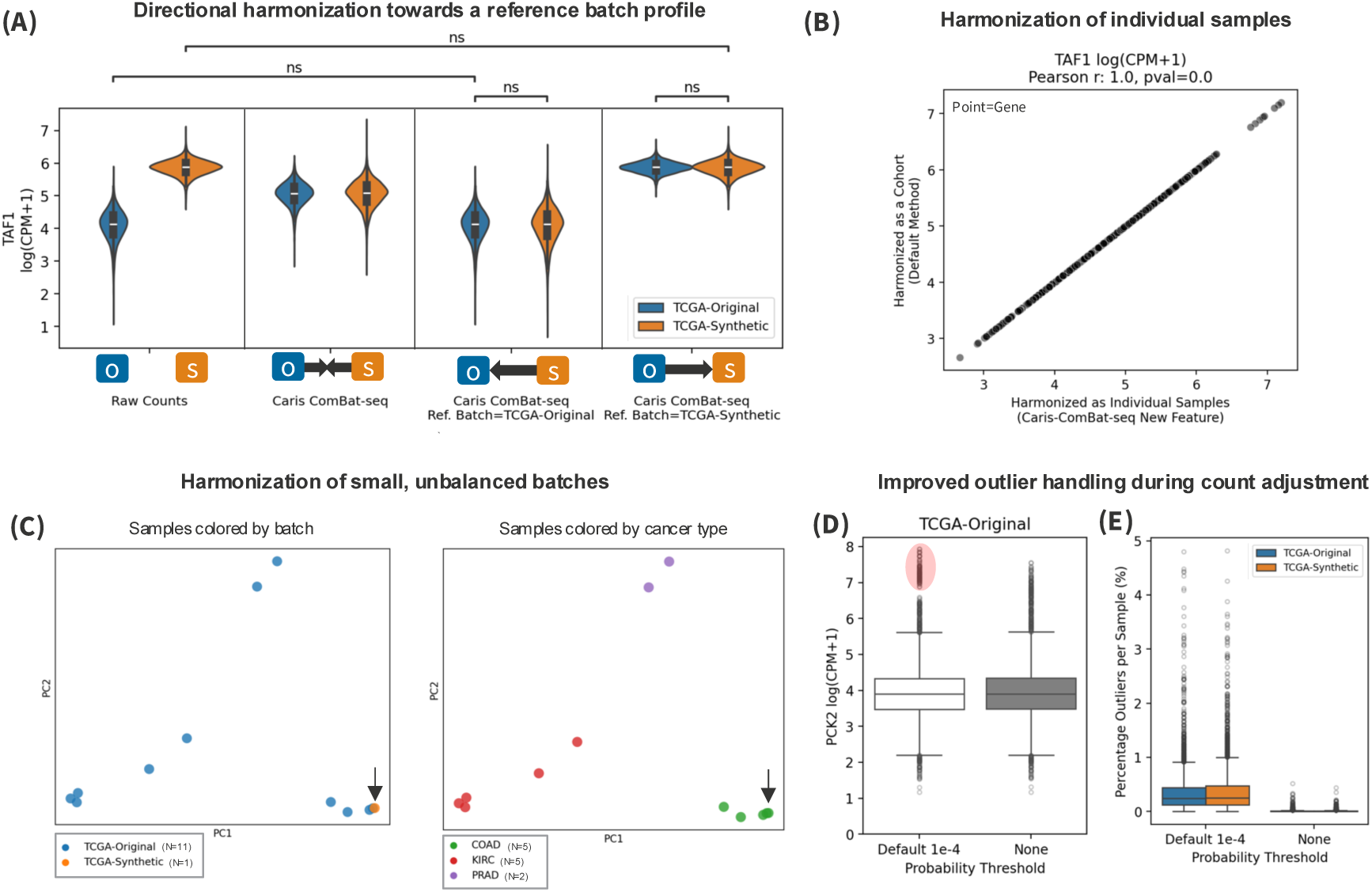
Caris-ComBat-seq implements three novel features to improve harmonized expression quality: directionally harmonizing towards a reference batch profile, harmonizing individual and small, unbalanced cohorts, and improved outlier count handling. **(A)** Violin plots of *TAF1* gene expression between batches (TCGA-Original: blue, TCGA-Synthetic: orange) in raw counts (left panel), adjusted counts harmonized with default Caris-ComBat-seq (middle left), adjusted counts harmonized with Caris ComBat-seq towards TCGA-Original as the reference batch (middle right), and adjusted counts harmonized with Caris ComBat-seq towards TCGA-Synthetic as the reference batch (right). Schematic with arrows and O/S boxes visually represent the directionality of adjustment applied to raw counts relative to batches. Statistical significance of mean expression differences between batches and harmonizing methods assessed with Mann-Whitney-Wilcoxon test two-sided with Benjamini-Hochberg (BH) correction. No significance (ns): 0.05 < *P* ≤ 1.0. **(B)** Correlation of per-sample adjusted *TAF1* expression when harmonized as a cohort (y-axis) or as individual samples (x-axis). **(C)** Sample-level (N = 12) principal component (PC) plots after Caris-ComBat-seq harmonized a small, batch-unbalanced cohort, colored by batch identity (left) and cancer type (right). COAD: colorectal adenocarcinoma, KIRC: kidney renal clear cell carcinoma, PRAD: prostate adenocarcinoma. **(D)** Boxplot of *PCK2* gene expression levels in TCGA-Original batch samples (N = 5,101) when harmonized using the pyComBat-seq default quantile mapping outlier probability threshold of 𝑝 < 10^−4^, compared to Caris-ComBat-seq without a probability threshold. Outlier counts are highlighted in red. **(E)** Percentage unadjusted outlier counts per sample in TCGA benchmarking dataset (N = 10,206) by batch (TCGA-Original: blue, TCGA-Synthetic: orange) when the default outlier probability threshold of 𝑝 < 10^−4^ is used compared to Caris-ComBat-seq’s new suggested threshold of no cutoff (”None”).

### Separating the training and adjusting steps

#### Harmonizing individual samples without having to re-harmonize entire cohorts

With the original implementations of ComBat-seq in both R and Python, as well as with ComBat-ref (21), a cohort would need to be re-harmonized every time a new sample is added, resulting in newly adjusted counts for *all* previous samples. Such a requirement is impractical in a clinical laboratory setting, where newly profiled samples must be continuously harmonized. Caris-ComBat-seq addresses this challenge by training a fixed model that learns batch effects across a representative set of samples, and then using this fixed model to adjust newly profiled samples. This approach reduces the computational and operational burden of the standard approach, because (1) the model does not have to be re-trained on a growing number of samples to allow adjusting newly profiled samples, and (2) previously profiled samples do not have to be re-adjusted using a continually re-trained model. To demonstrate the accuracy of this new approach to harmonizing individual samples, we show that samples harmonized with Caris-ComBat-seq as individual transcriptomes have identical count values compared to when harmonized in the traditional manner as part of a cohort (**Fig. 2B**; Pearson r = 1.0).

#### Harmonizing small, unbalanced batches

Traditionally, batches with few samples (i.e. <10) or very heterogenous samples are challenging to harmonize because it is hard to obtain accurate mean and dispersion estimates in these settings. Caris-ComBat-seq’s use of a fixed model trained on a large number of samples across batches overcomes this limitation by having robust, pre-calculated mean and dispersion estimates per gene per batch in the model. We demonstrate this capability with a toy dataset comprised of 12 samples: 11 from the TCGA-Original batch and one sample from the TCGA-Synthetic batch. The TCGA-Original batch contains samples from three cancer types – Kidney Renal Clear Cell Carcinoma (KIRC), Colorectal adenocarcinoma (COAD), and Prostate adenocarcinoma (PRAD) – while the single TCGA-Synthetic sample is a COAD type. Using the original R or Python implementation of ComBat-seq, this cohort would be impossible to harmonize because dispersion estimates can not be accurately made on a batch comprised of a single sample. However, with Caris-ComBat-seq we successfully harmonized the single TCGA-Synthetic sample batch and demonstrate that it correctly clusters together with COAD samples from the other batch (**Fig. 2C**).

### Improved outlier handling during quantile mapping improves harmonized expression quality

During the quantile mapping step used to adjust counts, the original ComBat-seq uses a probability threshold of 𝑝 < 10^−4^ to identify outlier counts that are not well-described by the modeled negative binomial distribution. If a count is identified as an outlier, it is not adjusted and remains as the original count value in the resulting adjusted counts matrix. This presence of original counts in harmonized expression matrices can create fictitiously small subpopulations of samples that appear to have ultra-high expression of a given gene, when really these are uncorrected counts of which the end-user is not aware. Caris-ComBat-seq implements two features to address this scenario: (i) the ability to set a custom probability threshold or entirely eliminate the threshold to adjust as many counts as possible, and (ii) the ability to return a data frame that flags which counts are deemed outliers during the quantile mapping step. Flagging outliers allows analysts to easily handle or exclude outliers from their desired analyses.

To illustrate the impact of unadjusted outliers, we selected a gene with a high outlier rate— *PCK2*—and compared expression of samples from the TCGA-Original batch when adjusted using an outlier probability threshold of 𝑝 < 10^−4^ (ComBat-seq default) versus not using a probability threshold (Caris-ComBat-seq recommendation). A small population of unadjusted counts, which manifest as high expressors, are visible when using an outlier threshold of 𝑝 < 10^−4^, but are successfully harmonized when the threshold is eliminated (**Fig. 2D**). Additionally, we see a higher proportion of unadjusted outlier counts per sample with the original outlier threshold, compared to when the threshold is eliminated (**Fig. 2E**).

### Parallelizing dispersion estimates and count harmonization accelerates run time

By separating the ComBat-seq framework into distinct training and adjusting steps, Caris-ComBat-seq can distribute batches and samples across compute resources to accelerate runtime compared to original implementations.

Batch correcting the TCGA benchmarking dataset of 10,206 samples and 9,932 genes takes 26.80 minutes to run with pyComBat-seq (**Fig. 3A**). To mirror the serial processing constraints of pyComBat-seq, we ran Caris-ComBat-seq on a single core and observed a similar wall time of 25.70 minutes (13.09 minutes of training + 12.61 minutes adjusting). However, when leveraging multiple cores to run Caris-ComBat-seq, we observed a marked ∼45% reduction in wall time to 11.71 minutes. Training wall time reduced by ∼40% to 7.92 minutes by parallelizing dispersion estimates per batch over independent cores, and harmonization wall time reduced by ∼70% to 3.79 minutes by distributing per sample harmonization over 16 cores in parallel. Chunking processes and optimizing the number of cores has the potential to further reduce wall time when running Caris-ComBat-seq.

**Figure 3.**
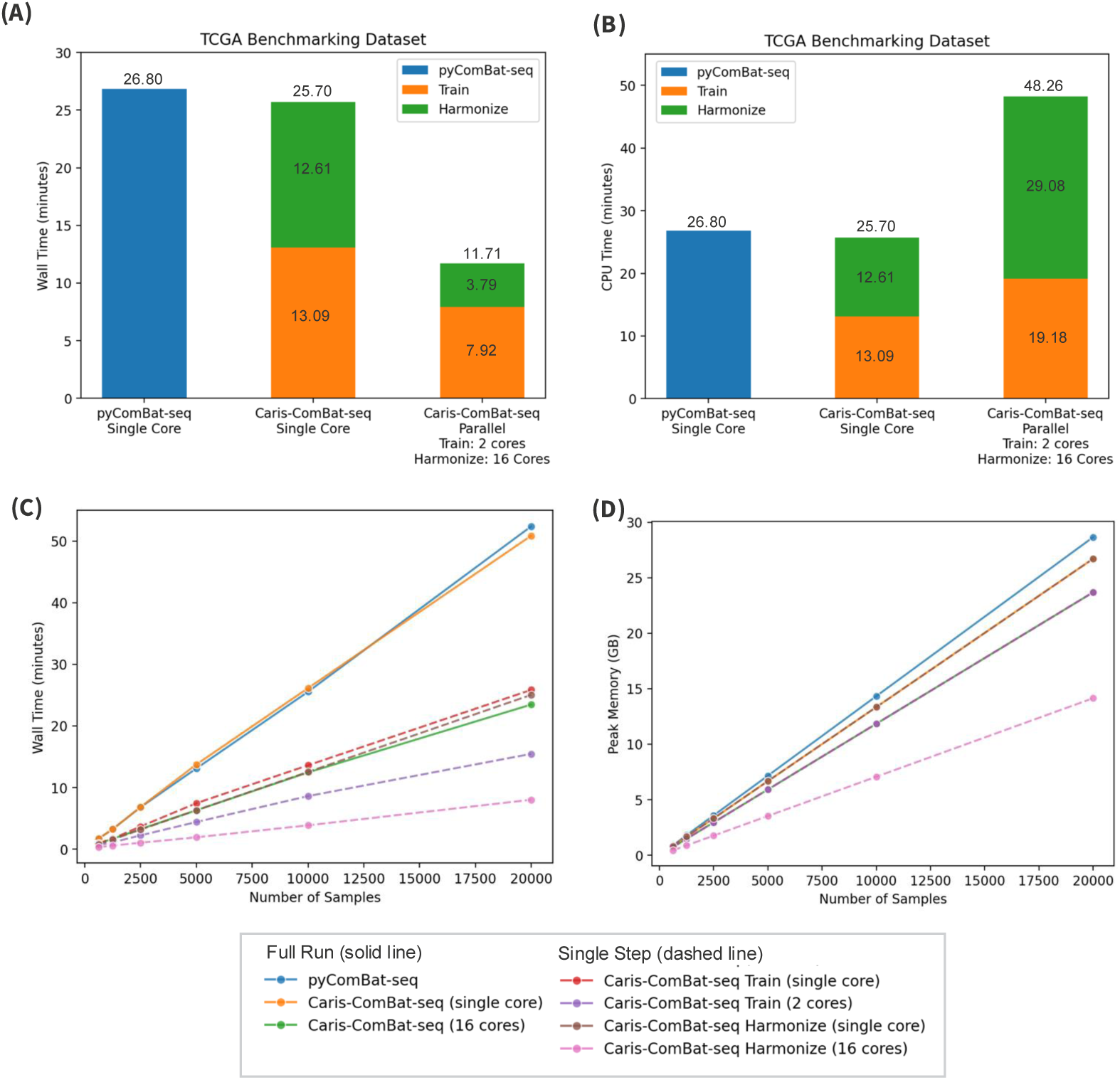
Caris-ComBat-seq runtime is greatly accelerated by parallelizing dispersion estimates (by batch) and count adjustment (by sample). (A-B) Comparison of **(A)** wall time or **(B)** CPU time (minutes; y-axis) across runs (x-axis) on TCGA benchmarking dataset comprised of 10,206 samples and 9,932 genes. Caris-ComBat-seq run time is visualized stacked by training and adjusting steps. Caris-ComBat-seq wall time is calculated when run on a single core, similar to pyComBat-seq, and when parallelizing the training (by batch) and adjusting (by sample) steps. Numbers above bars represent total wall time for run. Numbers on bars represent wall time for individual Caris-ComBat-seq steps. **(C-D)** Line plot depicting **(C)** wall time in minutes and **(D)** peak memory usage in GB across methods for subsamples of the TCGA benchmarking dataset varying from 625 to 20,000 samples. Runs include pyComBat-seq, as well as Caris-ComBat-seq training and adjusting steps implemented with single cores and parallelized across 16 cores.

We measured central processing unit (CPU) time to assess the total run time across resources with parallelization. When run in parallel across multiple cores, Caris-ComBat-seq has higher CPU time compared to both pyComBat-seq and Caris-ComBat-seq run on a single core (**Fig. 3B**). This result is expected as no modifications have been made to the underlying pyComBat-seq functions to increase their efficiency. Caris-ComBat-seq’s performance increases are enabled by the distribution of training and adjusting tasks across multiple resources to reduce wall time making it suitable to harmonize large expression cohorts of hundreds of thousands of samples given proper computational resources.

To compare the scalability of Caris-ComBat-seq to pyComBat-seq with cohort size, we ran both methods on cohorts comprised of 625, 1250, 2500, 5000, 10,000, and 20,000 samples from the TCGA benchmarking dataset and measured wall time and peak memory consumption **(Fig. 3C-D).** In cohorts where the number of samples exceeds the number of real samples in the TCGA dataset, we duplicated randomly selected samples to increase the cohort to the desired size. For both Caris-ComBat-seq and pyComBat-seq, wall time and CPU time scaled linearly as cohort size increased. By leveraging Caris-ComBat-seq’s ability to (i) harmonize newly profiled samples in parallel across 16 cores, and (ii) use a pre-trained model instead of recalculating the per-gene batch effect every time a new sample is added, the wall time to harmonize a new cohort of 20,000 samples is reduced by ∼85% (8.0min) in comparison to pyComBat-seq (52.4min) **(Fig. 3C).** Like the trends in wall time measurements, we also observed a markedly lower peak memory consumption when running the isolated harmonize step of Caris-ComBat-seq, compared to pyComBat-seq **(Fig. 3D).** These performance metrics highlight the benefits of separating the ComBat-seq statistical framework into distinct training and adjusting steps.

## DISCUSSION

The accurate, flexible, and efficient harmonizing of gene expression is critical for diverse research and clinical settings where different batches of RNA-seq data are analyzed together. Herein, we present Caris-ComBat-seq, which adds several new features to the ComBat-seq framework that make it better suited to a high-throughput clinical setting. Notably, the harmonized counts generated by Caris-ComBat-seq recapitulated those of the original ComBat-seq, demonstrating that our modifications did not impact the ability of the model to correct for batch effects while maintaining true biological heterogeneity. To maximize its utility, we have distributed Caris-ComBat-seq as an open-source python package with companion tutorial notebooks to help users adopt the tool (https://github.com/Caris-Life-Sciences/Caris-ComBat-seq). We have also made available all code and data necessary recreate the analyses in this manuscript (https://doi.org/10.5281/zenodo.17154014).

The first key advancement of Caris-ComBat-seq – directionally harmonizing towards a reference batch – enables selection of a high-quality “reference” batch that best recapitulates biological truth, similar to the recently developed approach taken by ComBat-ref (21). This modification potentially enables increased efficiency as well. First, since the counts for samples in the reference batch do not need to be adjusted, total run time may be reduced. Second, this feature could also reduce the ongoing harmonizing burden in a high-throughput laboratory setting by enabling a one-time adjustment of samples profiled with older platforms towards those in the batch profiled with the current platform. This approach would mean that newly profiled samples would not have to be harmonized; instead, older samples would be harmonized once to be compatible with newly profiled samples.

The second advancement of Caris-ComBat-seq is the flexible approach toward handling outliers in the data. Compared to ComBat-seq, where outlier counts with a probability threshold of 𝑝 < 10^−4^ are neither adjusted nor flagged (13), our method allows users to select if they want outliers flagged, or if they want the model to adjust them, with complete control over thresholding. In a clinical setting, this would enable a maximum number of samples to be harmonized and allow greater control and awareness over potential outliers.

The final and most important advancement of Caris-ComBat-seq—splitting the training and adjusting steps— eliminates the need for re-training the model with the addition of newly profiled samples, as long as they are consistent with the samples in the training set (i.e., sequenced using the same platforms and belonging to the same batches). This method also allows harmonizing small or unbalanced batches that are often encountered in real-world clinical trial cohorts, as we demonstrated by accurately clustering a single COAD sample that was harmonized with 11 other samples of different cancer types. Furthermore, this approach avoids having to run a memory- and time-intensive training step for every newly profiled sample that is added and enables parallelizing the adjusting step across multiple cores. Together, these scalability features are crucial in a molecular diagnostic setting where hundreds of samples are profiled each day.

To demonstrate the scalability and robustness of Caris-ComBat-seq in a real-world setting, we harmonized RNA-seq data from 383,331 clinical tumor samples profiled across two clinical assays: a legacy assay and Caris’ current Caris MI Tumor Seek Hybrid^™^. Using Metaflow to orchestrate our workflow and computational resources, we harmonized all samples in 1 hour and 3 minutes by paralleling in chunks of ∼385 samples using Amazon Web Services (AWS) batch. The resulting harmonized dataset showed a high-level of mixing between samples across assays, indicating that Caris-ComBat-seq successfully removed technical, protocol-driven batch effects from the RNA-seq dataset **(Figure S2.)** We have since adopted Caris-ComBat-seq as our production solution to harmonize RNA-seq count data from new samples profiled daily at Caris Life Sciences.

## CONCLUSION

In summary, we presented Caris-ComBat-seq, a variant of the ComBat-seq tool for harmonizing count data that offers key improvements in efficiency and harmonized data quality. Separating the training and adjusting steps, and directionally adjusting the data toward a high quality “reference” dataset allows newly profiled samples to be harmonized in a time-efficient manner – removing technical batch effects while retaining true biological variability. Improved management of outliers also gives users more control over the harmonizing process so that it meets their specific needs. We demonstrated the practical benefits of these improvements for a high-throughput setting, by using Caris-ComBat-seq to harmonize nearly half a million tumor samples analyzed by two different NGS platforms in a clinical diagnostic laboratory.

## AVAILABILITY & REQUIREMENTS

Project Name: Caris-ComBat-seq

Project home page: https://github.com/Caris-Life-Sciences/Caris-ComBat-seq and https://doi.org/10.5281/zenodo.17154014

Operating system: Platform independent Programming Language: Python

Other Requirements: inmoose, patsy, datetime, numpy, pandas, scipy, tqdm License: GNU GPL3

Any restrictions to use by non-academics: None

## LIST OF ABBREVIATIONS

AWS: Amazon Web Services
BH: Benjamini-Hochberg
BLCA: Bladder Cancer
BRCA: Breast Invasive Carcinoma
CESC: Cervical squamous cell carcinoma and endocervical adenocarcinoma
COAD: Colon adenocarcinoma
CPM: Counts per million
CPU: Central processing unit
ESCA: Esophageal carcinoma
GB: Gigabyte
GBM: Glioblastoma
HNSC: Head and Neck squamous cell carcinoma
InMoose: Integrated Multi Omic Open Source Environment
KIRC: Kidney renal clear cell carcinoma
KIRP: Kidney renal papillary cell carcinoma
LGG: Brain Lower Grade Glioma
LIHC: Liver hepatocellular carcinoma
LUAD: Lung adenocarcinoma
LUSC: Lung squamous cell carcinoma
NB: Negative binomial
NGS: Next-generation sequencing
ns: Not significant
OV: Ovarian serous cystadenocarcinoma
PAAD: Pancreatic adenocarcinoma
PC: Principal Component
PCA: Principal Component Analysis
*PCK2*: Phosphoenolpyruvate Carboxykinase 2
PRAD: Prostate adenocarcinoma
READ: Rectum adenocarcinoma
RNA: Ribonucleic Acid
RNA-seq: RNA Sequencing
SARC: Sarcoma
SKCM: Skin Cutaneous Melanoma
*SPEN*: Spen Family Transcriptional Repressor
STAD: Stomach adenocarcinoma
*t*-SNE: t-distributed stochastic neighbor embedding
*TAF1*: TATA-Box Binding Protein Associated Factor 1
TCGA: The Cancer Genome Atlas
THCA: Thyroid carcinoma
UCEC: Uterine Corpus Endometrial Carcinoma

## ACKNOWLEDGEMENTS

We thank Drs. Brady Gilg and Christian Frech, as well as Caris Life Science’s Data Science and Engineering Teams for their insights, constructive feedback, and continued support conceptualizing, refining, validating and productionizing Caris-ComBat-seq. We also thank Dr. Jennifer Ribeiro from the Scientific Writing team at Caris Life Sciences for valuable contributions to the preparation of this manuscript. Many of the results published here are based upon data generated by the TCGA Research Network: https://www.cancer.gov/tcga.

## AUTHOR INFORMATION

### Contributions

LMR performed all experiments and generated all figures in this manuscript. LMR and FPD designed, implemented and benchmarked Caris-ComBat-seq. LMR and FP wrote this manuscript with feedback from all co-authors. MV created production infrastructure to run Caris-ComBat-seq at scale. NSp created infrastructure to process and visualize Caris’ harmonized collection. LMR, MV, NSp, NSt and FPD reviewed code, manuscript text and provided significant scientific guidance. All authors read and approved the final manuscript.

## ETHICS DECLARATIONS

### Ethics approval and consent to participate

Not applicable.

### Consent for publication

Not applicable.

---

**Figure S1.**
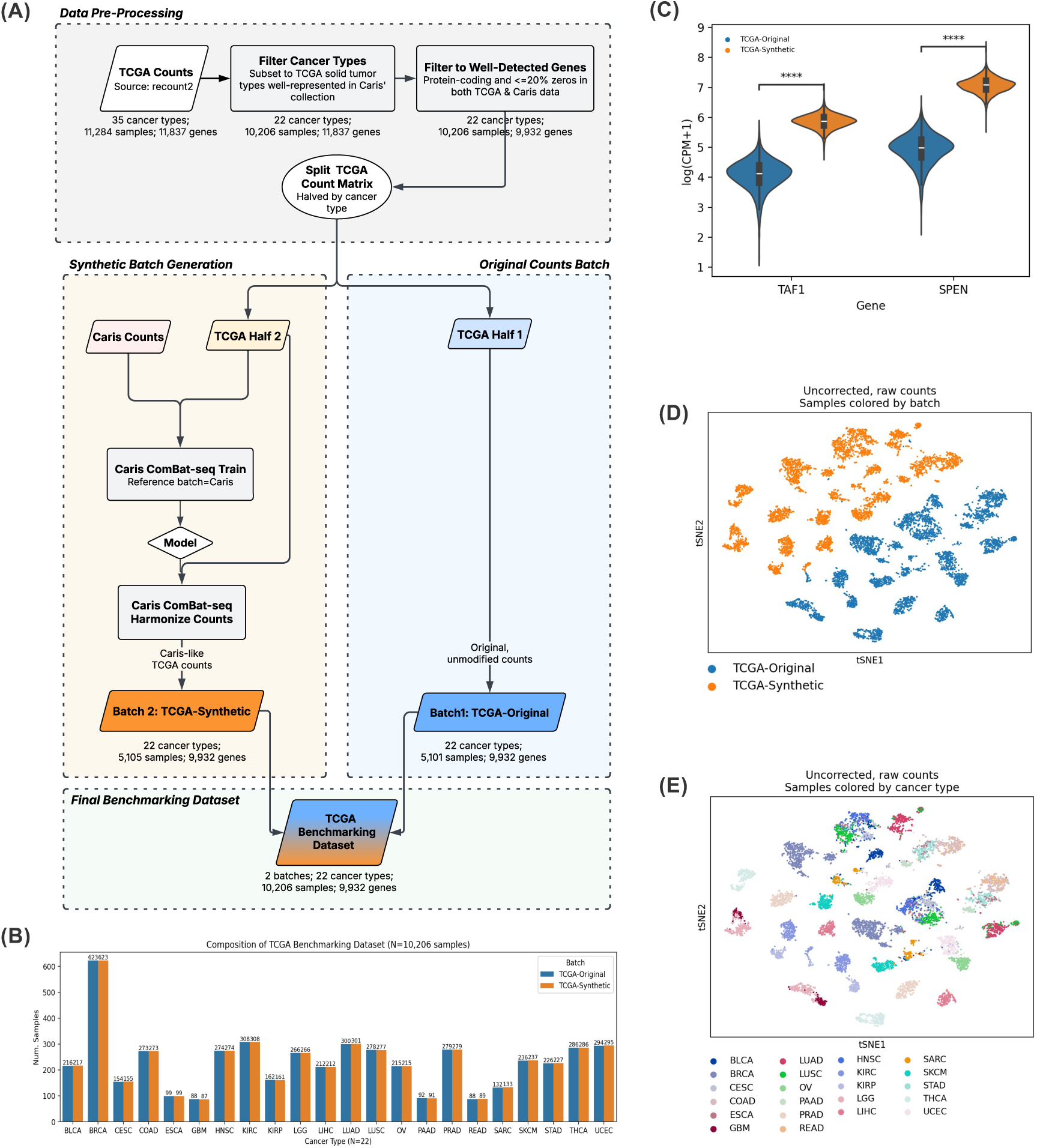
Generating a synthetic batch within TCGA for benchmarking harmonized expression quality on a real-world dataset of real-world. **(A)** Schematic depicting data pre-processing and use of Caris-ComBat-seq with directional reference batch harmonization to create a Caris-like synthetic batch within TCGA for benchmarking exercises. **(B)** Grouped bar plot depicting the sample numbers (y-axis) per cancer type (x-axis) in the TCGA-Original (blue) and TCGA-Synthetic (orange) batches. Cancer types are ordered alphabetically. Numbers above bars represent sample numbers per batch per cancer type. **(C)** Violin plot representing expression of two genes (*TAF1, SPEN*) with significantly different expression between TCGA-Original (N = 5,101 samples) and TCGA-Synthetic (N = 5,105 samples) batches. Statistical significance of mean expression differences between batches assessed with Mann-Whitney-Wilcoxon test two-sided with Benjamini-Hochberg (BH) correction. **** *P* ≤ 0.0001. (D-E) *t*-SNE visualization of sample-level transcriptome similarity and batch effect colored by **(D)** batch and **(E)** cancer type. Each point represents a sample (N = 10,206).

**Figure S2.**
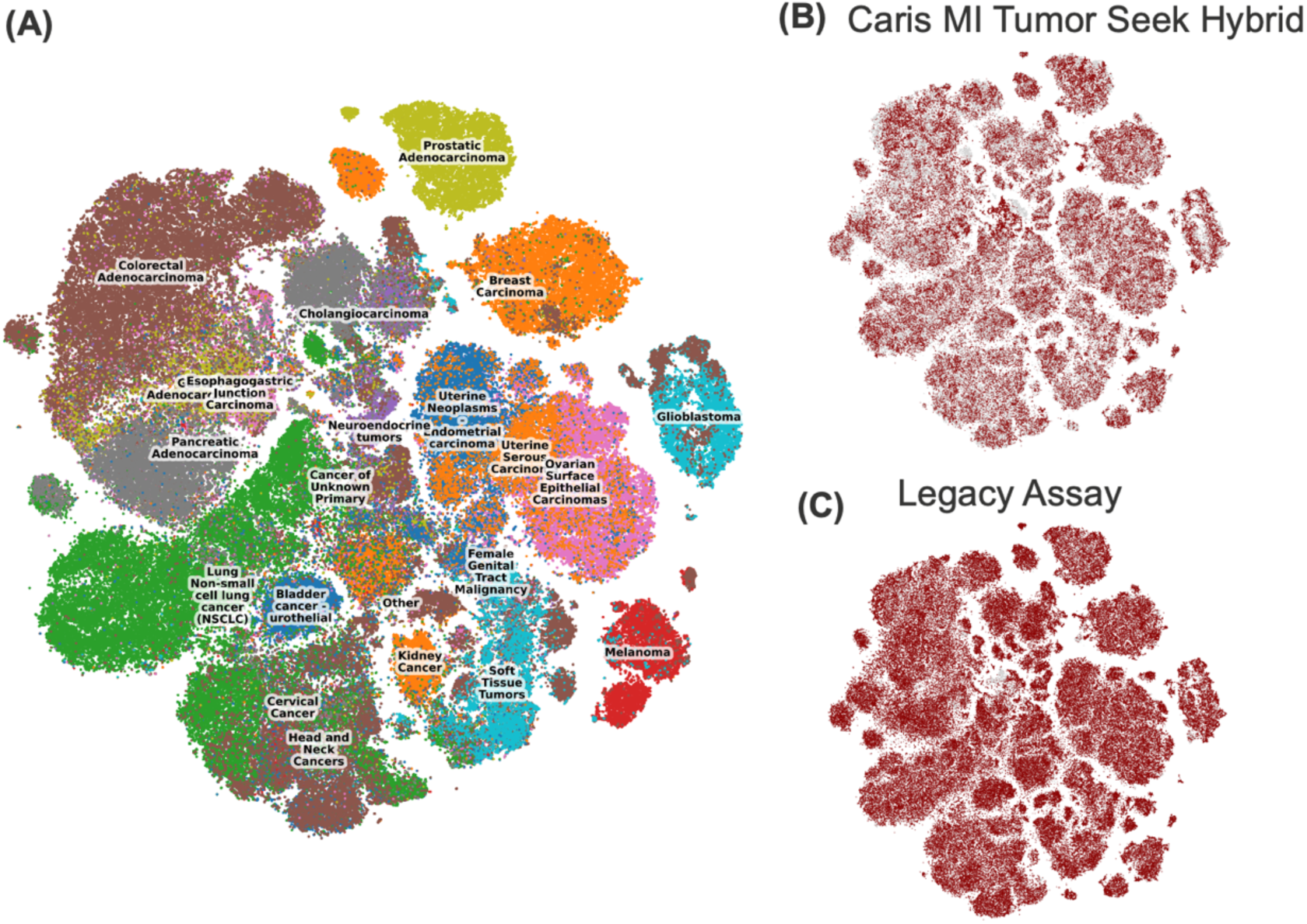
Harmonizing 383,331 clinical tumor samples with RNA-seq data across two clinical assays at Caris Life Sciences with Caris-ComBat-seq. (A-C) *t*-SNE visualization of Caris-ComBat-seq harmonized expression similarity and colored by **(A)** cancer type, and showing subsets of samples **(B)** profiled with the latest Caris MI Tumor Seek Hybrid (dark red) or **(C)** the legacy assay (dark red).

